# Opioids modulate Curiosity-Driven Exploration in Music

**DOI:** 10.64898/2026.05.05.722646

**Authors:** Claudia Alvarez-Martin, Raimund Buehler, Xim Cerda-Company, Gemma Cardona, Matthäus Willeit, Jacqueline Gottlieb, Giorgia Silani, Antoni Rodriguez-Fornells

**Author notes:** **Corresponding author:** Antoni Rodriguez-Fornells **Email:**. Both authors contributed equally to the present work (Claudia Alvarez-Martin, Raimund Buehler). Both authors declare co-senior contribution (Giorgia Silani, Antoni Rodriguez-Fornells). **Author Contributions: CAM:** Conceptualization; data curation; formal analysis; writing – original draft. **RB:** Conceptualization; data collection; writing – original draft. **XCC:** Conceptualization; formal analysis; writing – original draft. **GC:** Conceptualization; writing – original draft **MW:** Data collection. **JG:** Writing – review and editing. **GS:** Conceptualization; writing – original draft; funding acquisition; project administration; resources; supervision. **ARF:** Conceptualization; writing – original draft; funding acquisition; project administration; resources; supervision. **Competing Interest Statement:** The authors declare that they have no competing interests.

## Abstract

Curiosity, a key driver of exploration and learning, is reinforced by reward-related neurochemical systems, yet the role of the opioidergic system in modulating this behavior remains unclear. Music, as a highly rewarding stimulus, offers a unique context to investigate the neurochemical basis of curiosity, particularly the unexplored role of opioids in music-driven exploration. To fill this gap, we performed a double-blind within-subject pharmacological design, in which 26 participants received, in two different sessions, either a placebo or the opioid antagonist naltrexone. During each session, participants engaged in a music exploration/exploitation trade-off paradigm designed to assess their willingness to pay for exploring unfamiliar electronic music. Using logistic regression mixed-effects models, we found that while naltrexone did not affect overall curiosity ratings, it significantly reduced exploratory behavior in states of heightened curiosity. These findings suggest that the opioidergic system plays a critical role in regulating the relationship between curiosity and exploration, particularly in the context of novel and rewarding stimuli like music. Overall, the present research provides new and compelling evidence on the important relationship between curiosity and exploration and its regulation with the opioidergic neurotransmitter subsystem.

**Significance Statement:** The present research aimed to advance our understanding of the neurochemical mechanisms underlying curiosity and information seeking. In our study, we employed a pharmacological design to examine the role of the opioidergic system in music-related exploration. Using a novel music exploration/exploitation paradigm, we found that while naltrexone, an opioid antagonist, did not affect baseline curiosity ratings, it markedly reduced exploratory behavior during high-curiosity states in the presence of potential monetary losses. These results provide new evidence that opioidergic modulation plays a critical role in regulating curiosity-driven exploration. This new evidence might be relevant in the future for better understanding how neurochemical systems shape learning, motivation, and affective responses in complex cognitive domains such as music.

## Introduction

Curiosity, often defined as the intrinsic desire to seek out novel information or experiences, is a fundamental aspect of human cognition and behavior. Curiosity is not merely a passive state but an active drive that motivates individuals to explore their environment, acquire knowledge, and engage with unfamiliar stimuli. This drive is particularly pronounced when the stimuli are perceived as potentially rewarding or pleasurable, as curiosity often functions as a mechanism to reduce uncertainty and enhance learning (Berlyne, 1960; Loewenstein, 1994). Research has demonstrated that curiosity-driven exploration is closely tied to the anticipation of reward, which reinforces non-instrumental information-seeking activities and motivated learning (Blanchard et al., 2015; Cardona et al., 2026; Deaner et al., 2005).

From a neuropharmacological perspective, curiosity and information-seeking behaviors are underpinned by complex interactions within the brain’s reward systems. Studies have shown that curiosity elicits activation of reward-related dopaminergic pathways, particularly in regions such as the ventral striatum and prefrontal cortex (Bromberg-Martin & Hikosaka, 2009; Bromberg-Martin & Monosov, 2020; Gruber et al., 2014). These pathways are thought to play a critical role in reinforcing the pursuit of novel information by associating it with positive affect and motivational salience. Beyond dopamine, other neurotransmitter systems, such as the opioidergic system, have also been implicated in modulating the hedonic and motivational aspects of curiosity. For instance, research in animals has shown that blockading opioid receptors can diminish exploratory behavior and reduce the rewarding properties of novel stimuli, suggesting that the opioidergic system may influence the anticipatory rewarding value of curiosity-driven exploration (File, 1980; Katz & Gelbart, 1978; Achterberg et al., 2019). However, the exact role of opioids in curiosity and information-seeking remains underexplored, particularly in humans, where pharmacological interventions have yet to be systematically applied to study these phenomena.

Music, as a highly rewarding stimulus, provides a unique context for investigating the interplay between curiosity, reward, and neuropharmacology. Music is widely recognized for its profound impact on emotions and its ability to modulate affective states, making it one of the most universally rewarding human experiences. Recent studies have highlighted the role of the dopaminergic system in music-evoked pleasure, with neuroimaging and psychopharmacological research demonstrating its involvement in hedonic responses to music (Ferreri et al., 2019; Levitin & Menon, 2003; Salimpoor et al., 2011). Additionally, other neurotransmitter systems, including the opioidergic and oxytocinergic systems, have been suggested to contribute to the pleasure derived from music and its role in social bonding (Harvey, 2020; Savage et al., 2020). Despite this, the role of the opioidergic system in music reward remains somewhat ambiguous, with contradictory findings regarding its influence on pleasure and motivation (Laeng et al., 2021; Mas-Herrero et al., 2023; Goldstein, 1980; Mallik et al., 2017). This complexity makes music an ideal stimulus for exploring the neuropharmacological mechanisms underlying curiosity and reward, particularly in the context of novel or unfamiliar music, where curiosity-driven exploration may play a pivotal role.

In light of these considerations, the present research aims to bridge the gap in understanding the role of the opioidergic system in curiosity-driven exploration, specifically in the domain of music. By employing a double-blind, placebo-controlled, within-subject design and a novel exploration/exploitation paradigm, modelling participants’ willingness to engage with unfamiliar electronic music (Cardona et al., 2026), this study seeks to provide evidence of how pharmacological manipulation of opioid transmission (via administration of the unspecific opioid antagonist naltrexone), affects music curiosity and exploration in humans.

## Results

### Modelling participants’ choices: the effect of curiosity and reward

During the task, participants (n=26, 11 women, mean age 28.62 (sd = 11.39) years) were exposed to 10 s of a novel electronic song, after which they rated their level of curiosity using a 7-point Likert scale, and made a binary choice to continue listening to the song (“explore”) or switch to a familiar, favourite song (“exploit”). Each option was associated with a monetary reward (see Fig. 1). The psychometric curves shown in Fig. 2A plot the fraction of explore choices as a function of the reward difference between the two options (ΔMR), at different levels of curiosity.

**Figure 1.**
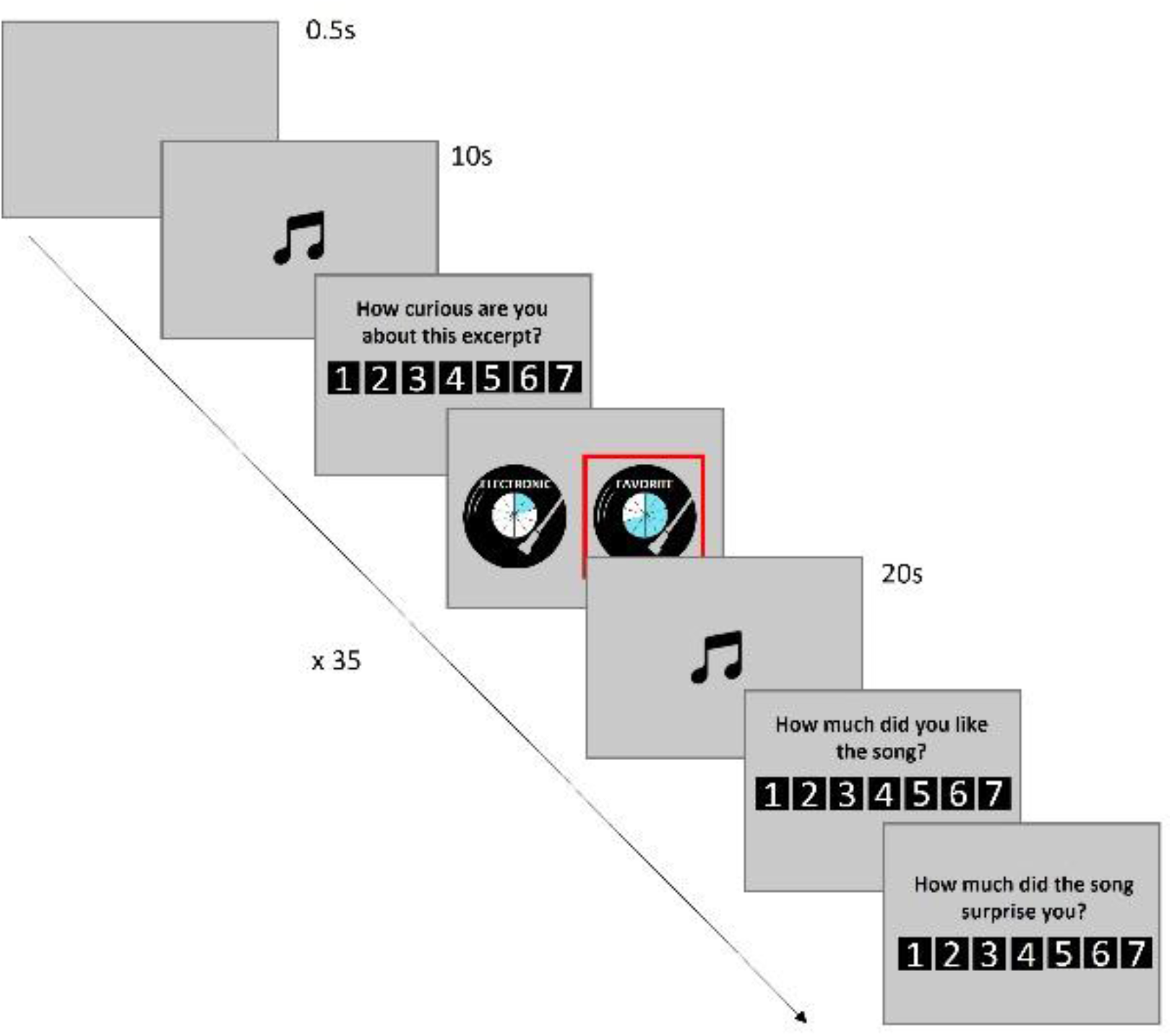
Schematic representation of the task structure. Participants listened to a preview of an unfamiliar novel song (10 s) and rated the elicited curiosity. Then, they had to choose between listening to the continuation of the track (LP with the word “electronic” written on it; novel option), or listening to an excerpt of a favourite song from a list they provided (LP with the word “favourite” written on it; favourite option). Each option was associated with a monetary reward represented by a 10-slices pie chart in the middle of the LP (one coloured slice = one euro). After the choice, music played for 40 s while a black musical note was displayed on the screen. In this example, the monetary reward for exploiting the favourite song is 7€, while the monetary reward for exploring the new song is 2€ (ΔMR = -5). Adapted from Cardona et al., 2026.

**Figure 2.**
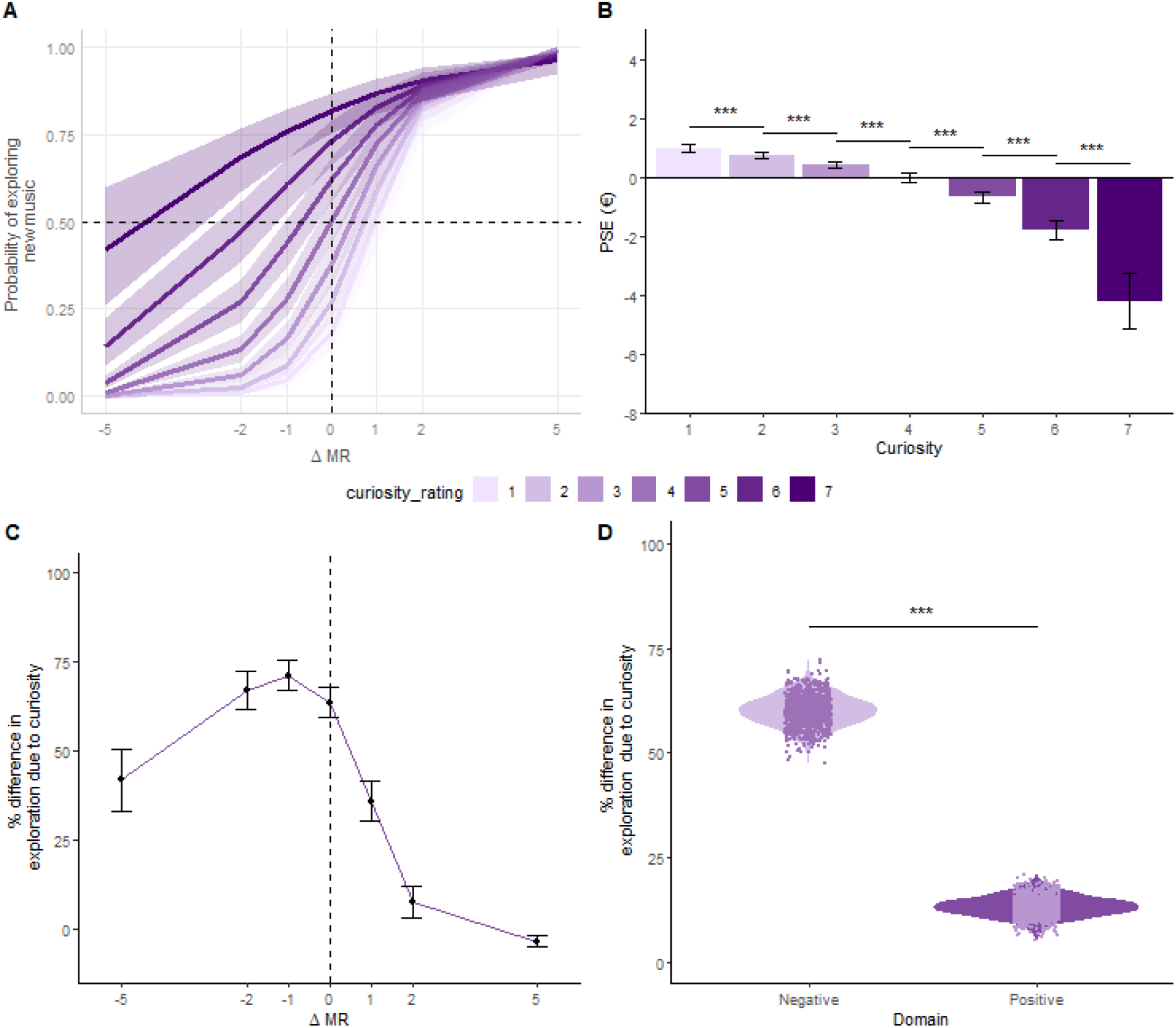
Representation of the interaction between curiosity and differential monetary value (ΔMR) and their effects on music choice. **A**. Psychometric curves corresponding to each possible curiosity rating reconstructed across participants (shadowed areas represent 95% confidence intervals). Notice the strong impact of curiosity specially in the negative domain. **B**. PSE (point of subjective equality) for each level of curiosity. Positive values indicate that participants would have to receive money in order to explore, and negative values indicate how much money participants would be willing to spend on exploring (the cost of a high curiosity state). Black bars represent standard error of the mean (SEM). Asterisks represent size effect reflected by the Cohen’s d (* = small effect (d ≥ 0.2), ** = moderate effect (d ≥ 0.5), *** = large effect (d ≥ 0.8)). **C**. Curiosity gradient. Differences in probability of exploring due to curiosity. For each differential value, the differences in probability between curiosity ratings of 1 and 7 are calculated. Error bars represent SEM. Note that the curiosity gradient is intensified in the negative domain, being maximum at -1 €. **D**. Mean curiosity gradient for the negative and positive domains.

Logistic models revealed significant effects of both predictors. The probability of choosing the new song increased as a function of its monetary reward (χ^2^(1)=182.53, p-value < 0.001) as well as its curiosity ratings (χ^2^(1)=105.31, p-value < 0.001, Fig. 2A and Table S1). Consequently, the PSE (point of subjective equality) decreased in function of curiosity – from +4.2 € for the highest curiosity rating to -1 € for the lowest rating (showing that participants were willing to pay approximately +4.2 € to listen to the new song and 1 € to listen to the favourite song).

The model also revealed a significant interaction between ΔMR and Curiosity (χ^2^(1)=61.20, p-value < 0.001, Table S2), whereby the impact of Curiosity was higher at negative versus positive ΔMR (Fig. 2A). As shown in Fig. 2C, the *curiosity gradient* (i.e., the difference in probability of exploring between curiosity levels of 7 and 1) peaked at ΔMR of -1 € (Fig. 2C), and was significantly stronger across all negative versus positive ΔMR levels (Fig. 2D). Thus, while participants tended to choose the song with the higher monetary reward, they were willing to sacrifice rewards to hear new songs, in proportion to their curiosity for the song.

### Effects of opioid receptor blockade on the modulation of curiosity and reward effects

To explore how pharmacological treatment affects exploration, we administered Naltrexone, an opioid antagonist, or a Placebo, within participants in two separate sessions. Logistic model fits revealed a three-way interaction between treatment, curiosity and ΔMR, whereby Naltrexone decreased the effect of curiosity for negative values of ΔMR – i.e., when participants had to pay to hear the novel song (χ^2^(1)=9.96, p-value=0.002; Table S2). This interaction can be seen in the lower levels of the psychometric curves under Naltrexone vs placebo at negative ΔMR (Fig. 3A). It is also expressed as a reduction in the curiosity gradient at negative ΔMR under Naltrexone – over all negative and positive ΔMR levels (Fig. 3C) and for each ΔMR level (Fig. 3B). Finally, comparison of the PSE values shows that, while Naltrexone did not significantly affect the amounts that participants paid for the low curiosity songs (curiosity = 1; 0.96 euro under placebo vs 1.02 euro after Naltrexone), it significantly reduced the willingness to pay for the high curiosity ones (curiosity = 7; – 5.82 euro for the novel song under placebo but only -3.56 Euros after Natlrexone (Fig. 3D). Importantly, Naltrexone did not impact the overall fraction of explore choices (Fig. 3E; χ^2^(1)=0.08, p =0.78, Table S3) or curiosity ratings (Fig. 3F; χ^2^(1)=1.44, p =0.23, Table S4). Thus, Naltrexone very specifically reduced the willingness to pay for novel songs in the absence of generalized effects on curiosity or the willingness to explore.

**Figure 3.**
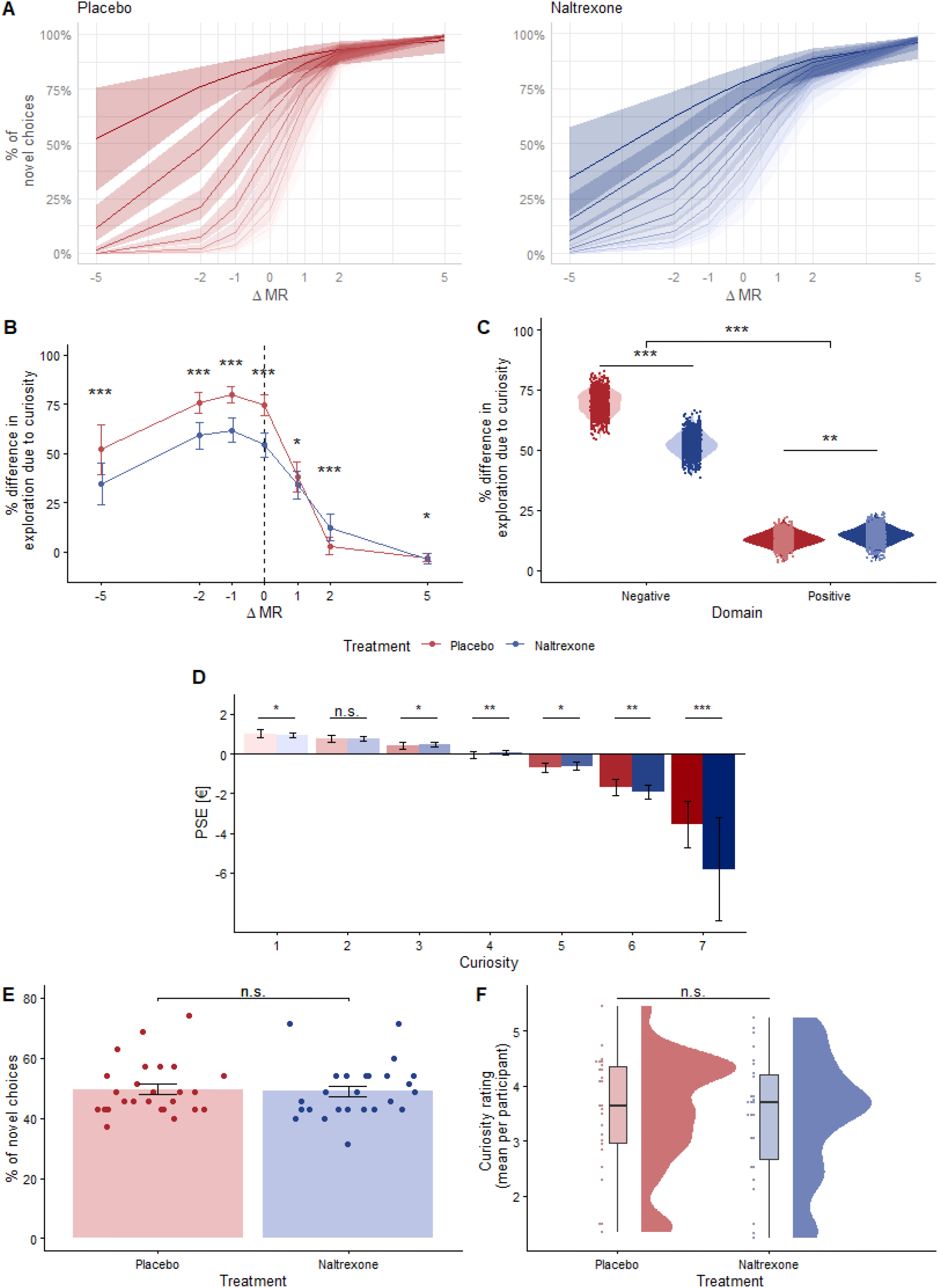
Representation of the interaction between curiosity and differential monetary value (ΔMR) and their effects on music choice for Placebo (red) and Naltrexone (Blue) **A**. Psychometric curves corresponding to each possible curiosity rating reconstructed across all participants for each treatment (shadowed areas represent 95% confidence intervals). Notice the strong impact of curiosity in the negative domain for Placebo, which is reduced by the administration of Naltrexone. **B**. Curiosity gradient. Differences in probability of exploring due to curiosity. For each differential value, the differences in probability between curiosity ratings of 1 and 7 are calculated. Error bars represent SEM. Note that the curiosity gradient is intensified in the negative domain, being maximum at -1 €. **C**. Mean curiosity gradient for the Negative and Positive ΔMR domains. **D**. PSE (point of subjective equality) for each level of curiosity. Positive values indicate that participants would have to receive money in order to explore, and negative values indicate how much money participants would be willing to spend on exploring (the cost of a high curiosity state). Black bars represent standard error of the mean (SEM). In Naltrexone, black horizontal bars represent Placebo values to ease comparison between treatments. Asterisks represent effect size for treatment for each curiosity rating, calculated using Cohen’s d (* = small effect (d ≥ 0.2), ** = moderate effect (d ≥ 0.5), *** = large effect (d ≥ 0.8)). **E**. Proportion of novel choices during sessions under each of the treatments. Dots represent individual values. **F**. Curiosity ratings during each session, classified by treatment. Values used correspond to the mean curiosity ratings for each session.

### Effects of opioid receptor blockade on satisfaction

Having demonstrated a specific effect of Naltrexone on choices and prospective curiosity ratings, we next examined if Naltrexone affected retrospective ratings, i.e., how much participants were satisfied with their choice.

To investigate satisfaction ratings, we fitted a mixed-effects model with subject as a random intercept to account for repeated measures. The model contained the main effects of treatment, choice, and domain, and the interactions of treatment with domain and choice (see Table S5 for the model selection steps and Table S6 for final model details). The results clearly showed a reduction of satisfaction in novel choices compared to favourite ones (χ^2^(1)= 149.76, p-value<0.0001, Fig.4A). Interestingly, a significant effect of domain was observed (χ^2^(1)= 23.29, p-value<0.0001), showing higher satisfaction in the negative compared to the positive domain (β=-0.64, p-value<0.0001, Fig.4B). Although naltrexone only slightly reduced satisfaction relative to placebo (β=-0.20, p=0.034), it had significant interactions with both choice (χ^2^(1)=7.34, p-value=0.007) and domain (χ^2^(1)=9.23, p-value=0.002). Under naltrexone, a larger decrease in satisfaction was observed for novel choices (β=-0.57, p-value=0.007), while this reduction was attenuated in the positive domain (β=0.63, p-value=0.002). These results point towards a selective modulation of satisfaction under naltrexone.

**Figure 4.**
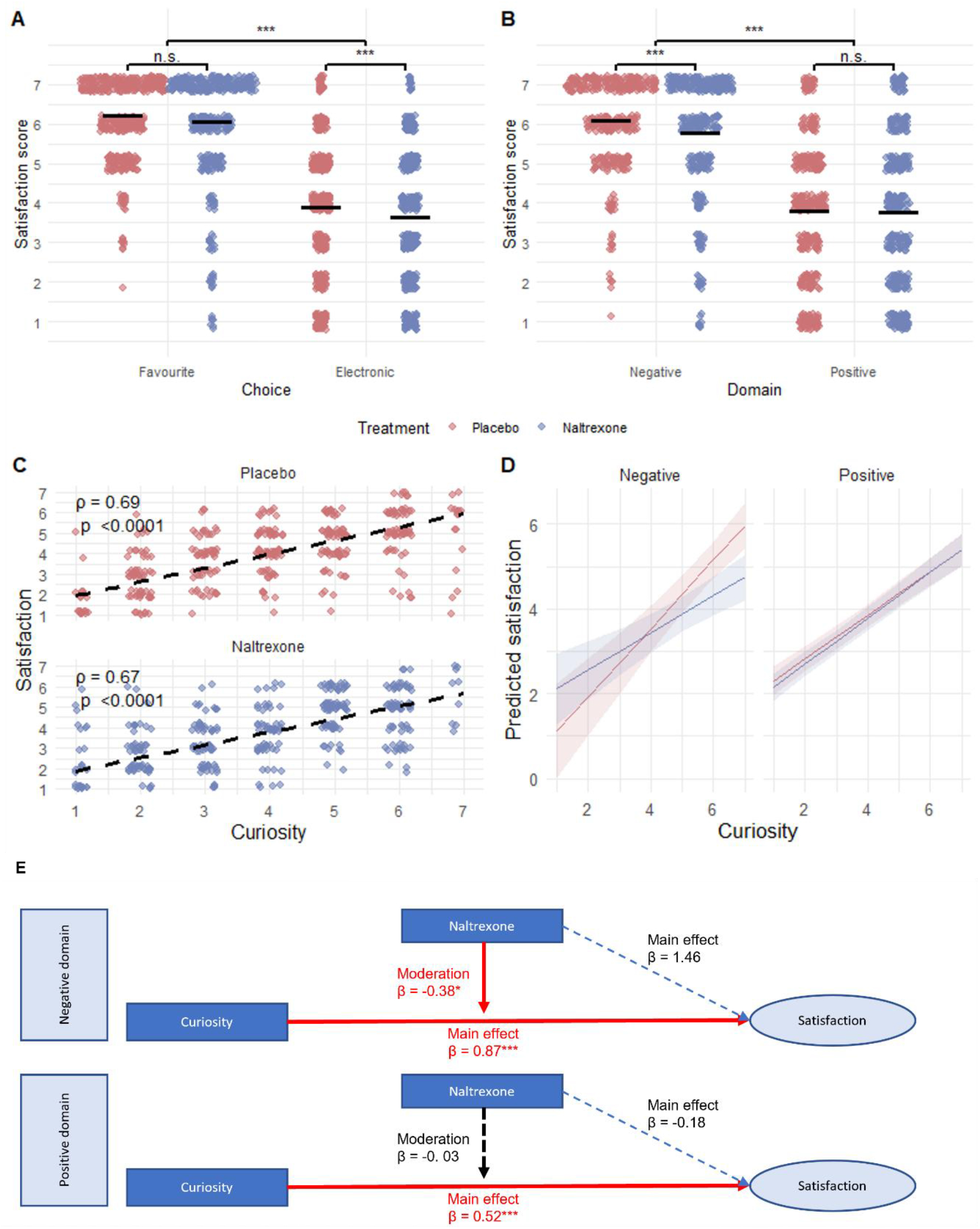
Impact of different variables on satisfaction ratings. **A**. Satisfaction ratings for favorite and novel options under each treatment. **B**. Satisfaction ratings for trials in each monetary domain. **C**. For novel choices only, correlation between curiosity and satisfaction under each treatment (using Spearman’s ρ). **D**. Predicted satisfaction for each curiosity level, for each treatment and domain. E. Schematic representation of the relationship between domain, treatment, and curiosity, and their effects on satisfaction for novel choices. The estimates were obtained by fitting a separate model for each monetary domain. As seen in the Figure, Naltrexone has a moderation effect on the relationship between curiosity and satisfaction only in the negative domain. Asterisks besides the β values indicate significance of the effect. Significance levels are indicated as follows: *** p < 0.001, ** p < 0.01, * p < 0.05, n.s. non-significant. For panels A and B, comparisons represent significance of the effects obtained from the lineal mixed-effects model reported in table S6.

To further study the effects of naltrexone on satisfaction ratings, we focused *exclusively on trials where participants chose to explore the novel option*, and therefore the satisfaction rating could reflect the “liking” of the continuation of the song. As expected, reported satisfaction after listening to the continuation of the novel songs correlated highly with the curiosity reported for the previews both under Placebo (ρ=0.69, p-value<0.0001) and under Naltrexone (ρ=0.67, p-value<0.0001, Fig. 4C). To explore the impact of treatment while considering the influence of curiosity on satisfaction for the novel songs, we fitted, only for novel choices, a new linear mixed-effects model that contained a three-way interaction between curiosity, domain, and treatment, all possible two-way interactions, and the main effects of the three variables (see Table S7 for model selection steps and Table S8 for mode). As expected, curiosity had a strong effect on satisfaction ratings (χ^2^(1)=382.64, p-value>0.0001). Overall, treatment with naltrexone did not show a significant main effect (χ^2^(1)=3.36, p-value=0.07). The three-way interaction had a significant effect (χ^2^(1)=6.28, p-value=0.012), indicating a moderation effect of naltrexone on the relationship between curiosity and satisfaction. As shown by the model predictions depicted in Figure 4D, in the positive domain the relationship between curiosity and satisfaction is maintained under placebo and naltrexone. In the negative domain, however, this relationship is weakened under naltrexone, where curiosity’s predictive power drops (Fig. 4E).

Overall, these results show that curiosity predicted satisfaction, which was reduced for novel songs. Importantly, while naltrexone had no effect on global satisfaction scores, it influenced satisfaction for novel songs in the negative ΔMR domain. Specifically, naltrexone moderated the relationship between curiosity and satisfaction: For novel songs, satisfaction was strongly predicted by curiosity, but this relationship was weakened under naltrexone in the loss domain.

## Discussion

Our study used a recently developed music curiosity trade-off task (Cardona et al., 2026) to investigate how the blockade of opioid transmission modulates curiosity and exploratory behaviors in humans. This task consisted of a trade-off which reflected the conflict between extrinsic reward (monetary reward), and the intrinsic reward associated with music listening. Following an exploration/exploitation design, participants had to choose between exploring novel, unfamiliar electronic music or exploiting their favorite songs, while the amount of monetary reward associated with the different options was manipulated.

Under placebo, the present study replicated previous results showing that participants’ decisions are influenced by both elicited music curiosity and monetary reward (Cardona et al., 2026). Notably, we observed a valence-dependent curiosity effect, showing that curiosity affects choices more strongly in the negative domain, where participants need to invest money to explore new songs, with participants becoming extremely selective and employing intrinsic curiosity signals to decide whether to explore. In other words, participants sacrificed money to explore only in those cases where curiosity was very high. In contrast, the effect of curiosity was drastically reduced in the monetary gain domain, where participants explored new pieces of music regardless of the elicited curiosity (probably reflecting a win-win scenario). In other words, when exploration is economically incentivized, participants opt to explore even when the song evoked little curiosity, likely prioritizing the extrinsic reward alongside the potential for new information (i.e., the new song). Thus, in high-curiosity states, the opportunity cost of losing money could be compensated by the anticipated pleasure derived from listening to the new song. It is interesting to note here that (i) a strong relationship was observed between curiosity states and satisfaction (a proxy to “liking”) after listening to the song, which confirms previous ideas on the strong relationship between reward anticipation and curiosity (Lowenstein, 1984), (see Fig. 4C) and (ii), satisfaction was indeed higher in the negative domain, when losing money was at stake (see Fig. 4b). This might also reflect that losing the opportunity of rehearing the new song had a larger impact than a potential monetary loss, pointing also to the characterization of curiosity as a loss-aversion phenomenon. This valence-dependent pattern of results is similar to previous studies showing that the decision to seek out new information is affected by the potential emotional value of receiving it (Charpentier et al., 2018).

The critical novel finding we observed concerns the effect of opioid receptor blockade. Administration of naltrexone affected neither music exploration (no reduction on the overall proportion of novel songs chosen) nor initial curiosity ratings (for both chosen and unchosen songs) (see Fig. 3e,f). However, a more detailed analysis revealed that under naltrexone, participants were markedly less likely to explore songs with high curiosity ratings when exploration required sacrificing money (negative domain). Considering that overall subjective curiosity was not affected, blockading opioid receptors acted selectively in the translation of curiosity into explorative behavior and decision making.

Importantly, our results are partially consistent with animal studies where an opioid antagonist administration also caused a reduction in exploratory behavior of a new environment (File, 1980; Katz & Gelbart, 1978) and playful behavior (Achterberg et al., 2019; Normansell & Panksepp, 1990; Trezza et al., 2010). Interestingly, a previous human study found a reduction of visual exploration (eye-face fixations) while participants viewed facial attractive photographs under naltrexone (Chelnokova et al., 2016). The authors concluded that opioid manipulation affects visual attention to socially relevant cues that might be important for obtaining potential rewarding information (Gottlieb, 2012; Maunsell, 2004) and for decision-making (Tatler et al., 2011). The present results add to current evidence in humans on the potential role of opioid regulation on reward motivation, specifically on the investment for salient rewarding conditions (Chelnokova et al., 2014; Eikemo et al., 2017; Korb et al., 2020; Weber et al., 2016, for evidence in non-human animals see Mahler & Berridge, 2012; Peciña & Berridge, 2013).

In order to explain this differential effect of naltrexone on information seeking and curiosity, it is important to analyze the type of computations in which participants engage to balance exploration-exploitation within the current paradigm. Participants might rely on (i) the potential value (anticipated pleasure) associated with listening to the pre-exposed new song (see Figure 4c for positive evidence on that), and (ii) the monetary opportunity cost. Considering that curiosity ratings and retrospective satisfaction scores (see Fig. 4a) did not differ due to naltrexone, we might assume that the anticipation of the rewarding value of the new song is preserved, including the savoring effects associated with high-curiosity states (Iigaya et al., 2016; Loewenstein, 1987; Zhu et al., 2017). This aligns with recent research indicating no direct effects of naltrexone in the modulation of anticipatory or subjective ratings of pleasure during music listening (Laeng et al., 2021; Mas-Herrero et al., 2023). Notice also that in our paradigm, not choosing to listen to the new electronic music piece, does not mean “not listening to music”: when participants disregard the novel option, they still hear a piece of music (a favorite song). Interestingly, van Steenbergen et al.(2019) (see also Eikemo et al., 2021) proposed that mu-opioid receptor stimulation might fine-tune reward valuation in affective contexts, especially for high-salience stimuli, rather than fully mediating hedonic states (see Buchel et al., 2018; Chelnokova et al., 2014; Eikemo et al., 2016; Petrovic et al., 2008). From this perspective, opioids might tune decision making by shifting this value function, in particular under loss or negative affect conditions. In this sense, opioid antagonists might dampen largely the potential subjective value for potential rewards affecting decision-making. In a similar vein, a recent study provided first evidence for the role of opioids in regulating approach motivation states, affecting attentional focus when anticipating a potential reward (van Steenbergen et al., 2025).

A detailed analysis of the satisfaction ratings for exclusively the selected novel songs revealed a strong correlation between the curiosity elicited by the novel excerpt and the satisfaction reported after listening to the continuation of the piece (Fig. 4c). Satisfaction ratings might reflect both the associated liking of the novel song and the potential impact of the economic reward associated with that choice. As commented above and similar to the results for overall curiosity, naltrexone revealed no significant effects on the global satisfaction ratings. However, a more detailed analysis of the novel choices revealed that opioid administration modulates the relationship between curiosity and satisfaction in the negative domain (see Fig. 4e): under naltrexone, the predictive properties of curiosity are weakened. This reduction in predictive power of curiosity could lead to the change in the decision-making patterns seen in the analysis of exploration: even when reporting high curiosity for a song, participants under naltrexone reduce their exploration when in the negative domain because the novel choices are less satisfying.

Another aspect to consider is that the value of the potential monetary loss could be more clearly affected under naltrexone, weighing more strongly in the decision to explore new songs with high curiosity ratings. For example, Petrovic and colleagues (2008) showed that an opioid antagonist (Naloxone) increased aversive processing of monetary losses, associated also with the amount of subjective unpleasantness. These results converge with previous studies pointing to the crucial role of opioids in aversive processing (Koppert et al., 2003, 2005; Narayanan et al., 2004; Skoubis et al., 2005; Wagner et al., 2007; Zubieta et al., 2001, 2003) suggesting that opioid blockade enhances aversive responses (Narayanan et al., 2004; Skoubis et al., 2005) and dysphoria. We could therefore reason that the processing of losses might have been amplified in our paradigm after naltrexone administration. In other words, increasing the subjective impact of the monetary loss could have made participants more cautious, inhibiting information seeking strategies.

It is important to also bear in mind that in the present paradigm, monetary reward is not immediately received by the participant; a time delay exists between the choice and the reception of the reward which does not correspond to the delay between the choice and listening to the song. Following this line of thought, one possible explanation for the reduction in information-seeking observed under high curiosity states could relate to reward impulsivity and delay discounting. Reward impulsivity is defined as the inability, in the face of a small immediate reward, to delay gratification and wait for a larger reward in the future. If we consider the decision to explore and being able to listen to high curiosity songs as an immediate reward (Ferreri et al., 2019) and the economic incentive as a delayed reward, our results could be explained by a reduction in reward impulsiveness under naltrexone administration, which would lead to the observed reduction of exploration of high curiosity choices in the negative domain. Indeed, naltrexone is used as a treatment for alcoholism, other addictions, and impulsive behavior disorders. However, direct evidence of the effect of naltrexone in reward impulsivity is scarce. A study by Kieres et al.(2004) showed that in rats, naltrexone was able to reduce the increase in delay discounting induced by Morphine administration, but it did not have an effect when administered alone. Moreover, to our knowledge only two studies directly investigated the effects of naltrexone in reward impulsivity in humans, with inconclusive results. Weber et al. (2016) found that naltrexone administration caused a non-significant reduction in reward impulsivity in a delay discounting task compared to placebo, while Mitchell et al. (2007) found the effects of naltrexone in reducing reward impulsivity to be personality-dependent. Taken together, the evidence regarding the involvement of the opioid system in reward impulsivity and delay discounting and its modulation by naltrexone may provide a partial explanation of the results observed in this study.

Furthermore, the mesolimbic opioid and dopamine systems are closely interconnected (Weber et al., 2016). Dopamine release in the nucleus accumbens (NAc) constitutes a critical step in the reward-seeking circuit, triggered by reward-inducing cues associated with music (Mavridis, 2015; Mueller et al., 2015). Opioids, however, can modulate dopamine release. Notably, naltrexone has been shown to block the alcohol- and feeding-induced dopamine release in the NAc (Benjamin et al., 1993; Taber et al., 1998), but not dopamine associated with monetary reward-predictive cues (Gowin et al., 2023). Further studies are necessary to understand how dopamine and opioid systems interact during our task and to disentangle the reduction of exploratory behavior induced by naltrexone.

Lastly, it is important to mention some limitations of the present study. First, our pharmacological manipulation used a non-specific opioid blocker (Naltrexone), preventing specific claims related to the mu-opioid system, as some of these effects could be mediated by kappa and, to a certain extent, delta receptors (Meier et al., 2021). Second, we selected new, unfamiliar songs that belonged to a particular genre, electronic music, which might have been affected by music preference. Further studies are needed to analyze information seeking related to the role of music genre preference and individual differences in music reward sensitivity.

To conclude, our results provide new insights on the involvement of opioidergic transmission in the processing of the rewarding properties of new music.

## Materials and Methods

### Participants

A total of 28 neurotypical participants took part in the study. Two participants did not understand the task and were removed from the analysis. The final sample consisted of 26 participants that completed both treatments (placebo and naltrexone, 11 women, mean age 28.62 +-11.39 years). Participants were included if they were at least 18 years old, fluent in German, had normal vision, and reported no history of psychiatric or neurological disorders or related medication use. Additional exclusion criteria included previous traumatic brain injury, substance (drug or alcohol) dependence, and any contraindications to naltrexone.

### Procedure

The experiment was conducted onsite at the University of Vienna in a within-subject design. During each session, participants received a pharmacological agent (see details in Pharmacological administration section) and performed a music curiosity trade-off task (detailed in Task section). This task took place among other non-related tasks (one eye tracking task and one reinforcement learning task), conducted on a laptop. Participants received a fixed compensation of 40€ per session, which was added to the variable monetary reward earned during this task. The study was approved by the ethics committee of the Medical University of Vienna (EK-1393/2017) in accordance with the principles of the Declaration of Helsinki, and participants signed an informed consent form.

### Pharmacological administration

This study is part of a larger protocol that employed a within-subject pharmacological design. Medications were supplied by the General Hospital of Vienna (AKH). Each participant experienced pharmacological conditions on separate testing days, with the order randomized for each participant. To ensure an adequate washout period, sessions were scheduled approximately two weeks apart (M = 15.8 days, SD = 7.44 days.).

Naltrexone (Dependex, 50 mg) was administered orally as a pill. In placebo conditions, participants received a pill with mannitol filler. Naltrexone is a nonspecific competitive opioid antagonist that preferentially blocks MOR, but also partially blocks kappa and delta receptors (Gonzalez & Brogden, 1988; Verebey et al., 1976). The dose of 50 mg was chosen based on previous studies, where naltrexone blocked 90% of mu-opioid receptors at this dose (Weerts et al., 2013) and produced minor side effects in healthy volunteers. Naltrexone reaches a peak plasma concentration at 1 h following oral administration (Gonzalez & Brogden, 1988; Verebey et al., 1976), which was accounted for in the timing of Naltrexone administration (Gonzalez & Brogden, 1988; Gossen et al., 2012). To check blinding success, participants guessed whether they had received the active drug or placebo at the end of each session. Participants were unable to reliably distinguish between treatments, confirming effective blinding (χ^2^(1) = 2.53, p = 0.112).

### Musical stimuli

Musical stimuli consisted of 140 unfamiliar electronic music tracks, from which excerpts of 30s were extracted. All the excerpts were split into a preview (the first 10 s) and a continuation (the next 20 s). Stimuli were selected by Indiemono^®^, a music curator company. Importantly, novel stimuli did not contain lyrics nor highly salient features (e.g. a single instrument leading the melody, highly noticeable audio distortions, sounds resembling human voices), and curators tried to avoid including multiple songs by the same artist. From the selected pieces, we used the MIR Toolbox for Matlab (Lartillot & Toiviainen, 2007) to calculate several properties such as loudness, percussive onsets, tempo, rhythmic complexity and musical contrast. Subsequently, the excerpts were divided into four balanced lists showing no significant differences regarding these metrics (ps > 0.6, ANOVA tests of unequal variances). These four lists were counterbalanced across subjects and sessions.

In addition, each participant was asked to provide a list including the 20 favourite or preferred songs they had listened to the most during the last year (Cardona et al., 2026). From each song of that list, the first 20 s were extracted. All stimuli were automatically normalized and faded (2.5 s in and 2.5 s out) using the *pydub* (Robert et al., 2018) library from Python. Stimuli were presented using headphones and participants were able to adjust the volume at the beginning of the task.

### Task design

Participants were exposed to 35 trials during each session. In each trial, participants listened to the preview of an novel music excerpt while a black note was displayed on screen (see paradigm in Fig. 1). Participants were then asked to rate their curiosity about the excerpt using a 7-point Likert scale (where 1 meant Not curious at all and 7 meant Extremely curious). Once they provided their rating, participants could choose between listening to the continuation of the novel excerpt (i.e., novel option) or to one of the songs included in their favourite playlist, chosen at random (i.e., favourite option). This choice was represented on screen by two long-play (LP) images with the labels “electronic” and “favourite” on top. The presentation side of the LPs was counterbalanced across participants. A 10-slices pie chart was displayed at the centre of each LP, with the number of coloured slices representing the economic reward associated with each of the options (ranging from one to ten euros). During each session a total of five trials corresponding to each differential value were presented pseudorandomly.

After the choice, participants listened to 20 s of music corresponding to the rest of the novel excerpt or the beginning of the favourite song, depending on the option selected. After listening to it, participants were asked to rate, using a 7-point Likert scale, their 1) satisfaction (i.e., “How much did you like the song?”, from 1=I didn’t like it at all, to 7=I liked it a lot); and 2) surprise (i.e., “How much did the song surprise you?”, from 1=No surprise at all, to 7=High surprise). Participants were instructed to report the absolute value of this rating, disregarding whether the surprise was positive (the continuation ended up being better than expected) or negative (they were disappointed by its unfolding).

Finally, at the end of each session two trials were pseudorandomly selected and participants received the money associated with the choice from these trials. The selection of the trials was manipulated to keep the final reward for each session between 2 and 10 euros. As indicated previously, this reward was added to the fixed amount that participants received for their participation. The task was presented using Psychopy version 2022.1.4 (Peirce et al., 2019).

### Data analysis

All analyses were performed using R (R Core Team, 2022) version 4.2.0, and generalized mixed-effect regression models were fitted using the lme4 package (Bates et al., 2015). Model selection steps and results were reported following the guidelines from Meteyard and Davies (2020).

#### Analysis of music choice

Based on the economic reward associated with each option (novel or favourite), we calculated the differential monetary reward value (ΔMR) as the difference between the electronic and favourite options. Specifically, ΔMR took values of {-5, -2, -1, 0, 1, 2, 5} euros, where negative values indicated a lower economic reward for the electronic option compared to the favourite, and positive values indicated the opposite.

To analyse the likelihood of exploring the novel option, we used generalized mixed-effects models (binomial family), with ‘subject’ as random effect. Based on previous research (Cardona et al., 2026), we defined a control model which included the main effects of ΔMR and curiosity, along with their interaction. Psychometric curves were predicted using the *ggeffects* package (Lüdecke, 2018) to depict the probability of exploration as a function of the ΔMR and the curiosity level. This control model served as a null model, providing a baseline to assess how the variable of interest of this study (namely, treatment) impacted exploratory music behaviour.

To further explore the interaction between curiosity and ΔMR, we computed the point of subjective equality (PSE) and the curiosity gradient. The former indicates the ΔMR value at which participants showed no preference between the novel and favourite options (i.e., exploration probability of 50%). We calculated the PSE for each curiosity rating using Equation 1, obtaining the values of ΔMR that would mean a 50% probability of choosing novel music. The coefficients used in these equations were obtained by generating 1000 simulations of each model with a bootstrapping approach, thus obtaining a distribution of PSE values and allowing for statistical comparisons. PSE comparisons between curiosity levels and treatments were performed by computing Cohen’s *d*, which allows for the determination of the effect size.

The curiosity gradient (probability of exploring a song with a curiosity score of 7 minus the probability of exploring with a curiosity of 1) represents how much a participant’s likelihood of exploring changes in response to curiosity at each ΔMR value. It captures the “sensitivity” of exploratory behaviour to different curiosity levels, by quantifying how exploration probability shifts as curiosity levels increase or decrease. For each participant, this gradient reflects the effect of curiosity on exploration across different ΔMR values. The curiosity gradient was also obtained using a bootstrapping approach, generating 1000 model simulations and obtaining from each of them the probability of exploration when curiosity was rated as a 1 or a 7 for each value of ΔMR. The difference in exploration between both curiosity levels was calculated and multiplied by 100 to obtain the percentage difference. We further summarized this variable by averaging, for each simulation, the difference in exploration by monetary domain (i.e., Negative when ΔMR < 0 and Positive when ΔMR > 0). The comparison between treatments of the curiosity gradient for each value of ΔMR and domain was performed by using Cohen’s *d* to assess the effect size.

To assess the effects of our variable of interest (treatment) on music exploration, we fitted a generalized mixed-effects model that incorporated the effect of treatment and its interaction with Curiosity and ΔMR, using the control model as a baseline. A backward approach was used, beginning with all possible main effects and interactions, which were sequentially removed based on their significance within the model. At each elimination step, model performance was compared with the control model using Akaike Information Criterion (AIC) and the Likelihood Ratio Test (LRT). Finally, the model with the lowest AIC score was selected as the final model. The same approach was used to generate the PSE and curiosity gradient for each treatment, in this case using Equation 2 to calculate the PSE values to take into consideration the effect of treatment.

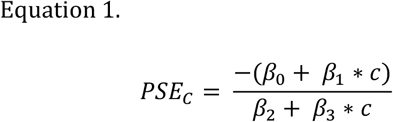

Where β_0_ represents the coefficient of the intercept, β_1_ represents the coefficient of curiosity, β_2_ represents the coefficient of ΔMR, β_3_ represents the coefficient of the interaction between curiosity and ΔMR, and *c* represents the curiosity level.

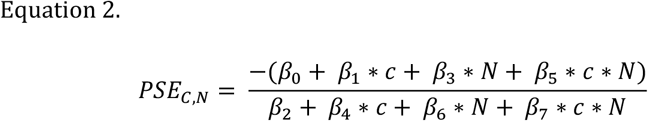

Where β_0_ represents the coefficient of the intercept, β_1_ represents the coefficient of curiosity, β_2_ represents the coefficient of ΔMR, β_3_ represents the coefficient of treatment, β_4_ represents the interaction between curiosity and ΔMR, β_5_ represents the interaction between curiosity and treatment, β_6_ represents the interaction between ΔMR and treatment, β_7_ represents the coefficient of the interaction between curiosity, ΔMR, and treatment, *c* represents the curiosity level (ranging from 1 to 7), and *N* represents naltrexone treatment (0 for placebo, 1 for naltrexone)

## Acknowledgments

ARF has been suported by the FIAS fellowship Program, co-funded by the European Commission, Marie-Skłodowska-Curie Actions - COFUND Program, Grant n°945408. CAM is supported by a predoctoral contract (FPI, PRE2022-104055) funded by the Agencia Estatal de Investigación (AEI). This project received funding from grants PID2023-151083NA-I00 funded by MICIU/AEI/10.13039/501100011033 and by ERDF/EU, grant PID2021-127130NB-100 funded by MICIU/AEI/10.13039/501100011033, and grant 10.55776/PAT3432624 funded by the Austrian Science Fund (FWF). We thank CERCA Programme / Generalitat de Catalunya for institutional support.

