## Supplemental tables for "Opioids modulate Curiosity-Driven Exploration in Music"

### Supplementary information

**Table S1.** Model details: interaction between curiosity and monetary reward on music choice.

| Fixed effects | | | | | |
|---|---|---|---|---|---|
| | Beta | SE | CI | z | p-value |
| Intercept | -1.99 | 0.23 | 0.4 - 0.6 | -8.77 | < 0.001 |
| Curiosity | 0.5 | 0.05 | 1.47 - 1.98 | 10.26 | < 0.001 |
| ΔMR | 1.72 | 0.13 | -0.24 - -0.15 | 13.51 | < 0.001 |
| Curiosity:ΔMR | -0.19 | 0.02 | 0.4 - 0.6 | -7.82 | < 0.001 |
| Random effects | | | | | |
| | Variance | sd | | | |
| Subject | 0.317 | 0.563 | | | |
| Model fit | | | | | |
| R2 | Marginal | Conditional | | | |
| | 0.76 | 0.78 | | | |

**Table S2.** Model details: effect of treatment on proportion of novel choices.

| Fixed effects | | | | | |
|---|---|---|---|---|---|
| | Beta | SE | CI | z | p-value |
| Intercept | -0.01 | 0.08 | -0.17 - 0.15 | -0.16 | 0.872 |
| Naltrexone | -0.03 | 0.09 | -0.21 - 0.16 | -0.28 | 0.777 |
| Random effects | | | | | |
| | Variance | sd | | | |
| Subject | 0.055 | 0.234 | | | |
| Model fit | | | | | |
| R2 | Marginal | Conditional | | | |
| | 0 | 0.02 | | | |

**Table S3.** Model details: effect of treatment on curiosity ratings for all novel songs.

| Fixed effects | | | | | |
|---|---|---|---|---|---|
| | Beta | SE | CI | z | p-value |
| Intercept | 0.02 | 0.12 | -0.2 - 0.25 | 0.2 | 0.843 |
| Naltrexone | -0.05 | 0.04 | -0.12 - 0.03 | -1.2 | 0.23 |
| Random effects | | | | | |
| | Variance | sd | | | |
| Subject | 0.33 | 0.574 | | | |
| Residual | 0.683 | 0.826 | | | |
| Model fit | | | | | |
| R2 | Marginal | Conditional | | | |
| | 0 | 0.33 | | | |

**Table S4.** Model details: effect of treatment on the interaction between curiosity and monetary reward.

| Fixed effects | | | | | |
|---|---|---|---|---|---|
| | Beta | SE | CI | z | p-value |
| Intercept | -2.67 | 0.34 | -3.32 - -2.01 | -7.96 | < 0.001 |
| Curiosity rating | 0.65 | 0.07 | 0.51 - 0.79 | 8.85 | < 0.001 |
| ΔMR | 2.41 | 0.23 | 1.96 - 2.86 | 10.57 | < 0.001 |
| Naltrexone | 1.06 | 0.38 | 0.32 - 1.8 | 2.82 | 0.005 |
| Curiosity: ΔMR | -0.29 | 0.04 | -0.37 - -0.21 | -7.1 | < 0.001 |
| Curiosity:Naltrexone | -0.24 | 0.09 | -0.41 - -0.07 | -2.74 | 0.006 |
| ΔMR:Naltrexone | -1.1 | 0.27 | -1.63 - -0.57 | -4.09 | < 0.001 |
| Curiosity:ΔMR:Naltrexone | 0.16 | 0.05 | 0.06 - 0.26 | 3.16 | 0.002 |
| Random effects | | | | | |
| | Variance | sd | | | |
| Subject | 0.324 | 0.569 | | | |
| Model fit | | | | | |
| R2 | Marginal | Conditional | | | |
| | 0.79 | 0.81 | | | |

**Table S5.** Model selection steps for the variables affecting satisfaction ratings

| Sampling units | N total observations = 1560<br>N subjects = 26 | | | | | | | | | |
|---|---|---|---|---|---|---|---|---|---|---|
| Model specification | Model name | Nested model | Fixed effects removed | Random effects | Model fit | | | LRT test against nested | | |
| | | | | Subjects | AIC | BIC | LL | Df | X2 | p-value |
| ttm:choice:domain + ttm:choice + choice:domain + ttm:domain + choice + ttm + domain | M0 | - | - | Intercept | 5280.5 | 5334 | -2630.3 | - | - | - |
| ttm:choice + choice:domain + ttm:domain + choice + ttm + domain | M1 | M0 | ttm:choice:domain | Intercept | 5279.3 | 5327.5 | -2630.7 | 1 | 0.79 | 0.373 |
| ttm:choice + ttm:domain + choice + ttm + domain | M2final | M1 | choice:domain | Intercept | 5280.4 | 5323.2 | -2632.2 | 1 | 3.10 | 0.078 |
| ttm:domain + choice + ttm + domain | M3 | M2final | ttm:choice | Intercept | 5285.8 | 5323.2 | -2635.9 | 1 | 7.35 | 0.007 |
| ttm:choice + choice + ttm + domain | M4 | M2final | ttm:domain | Intercept | 5287.6 | 5325.1 | -2636.8 | 1 | 9.24 | 0.002 |

**Table S6.** Model details: effect of treatment, choice, and domain on satisfaction scores

| Fixed effects | | | | | |
|---|---|---|---|---|---|
| | Beta | SE | CI | t | p-value |
| Intercept | 6.30 | 0.17 | 5.97 – 6.63 | 37.88 | <0.0001 |
| Novel choice | -1.89 | 0.15 | -2.20 - -1.59 | -12.24 | <0.0001 |
| Naltrexone | -0.2 | 0.09 | -0.39 - -0.01 | -2.13 | 0.034 |
| Positive domain | -0.74 | 0.15 | -1.04 - -0.44 | -4.83 | <0.0001 |
| Naltrexone:Novel choice | -0.57 | 0.21 | -0.98 - -0.16 | -2.71 | 0.007 |
| Naltrexone:Positive domain | 0.63 | 0.21 | 0.23 – 1.04 | 3.04 | 0.002 |
| Random effects | | | | | |
| | Variance | sd | | | |
| Subject | 0.60 | 0.78 | | | |
| Model fit | | | | | |
| R2 | Marginal | Conditional | | | |
| | 0.42 | 0.58 | | | |

**Table S7.** Model selection steps for the variables affecting satisfaction ratings only in novel choices.

| Sampling units | | N total observations = 812<br>N subjects = 26 | | | | | | | | |
|---|---|---|---|---|---|---|---|---|---|---|
| Model specification | Model name | Nested model | Fixed effects removed | Random effects | Model fit | | | LRT test against nested | | |
| | | | | Subjects | AIC | BIC | LL | Df | X2 | p-value |
| ttm:domain:curiosity + ttm:domain + ttm:curiosity + domain:curiosity + ttm + domain + curiosity | M0final | - | - | Intercept | 2585 | 2631.9 | -1282.5 | - | - | - |
| ttm:domain + ttm:curiosity + domain:curiosity + ttm + domain + curiosity | M1 | M0final | ttm:domain:curiosity | Intercept | 2589.3 | 2631.6 | -1285.6 | 1 | 6.32 | 0.012 |

**Table S8.** Model details: effect of treatment, curiosity, and domain on satisfaction scores only for novel choices.

| Fixed effects | | | | | |
|---|---|---|---|---|---|
| | Beta | SE | CI | t | p-value |
| Intercept | 0.318 | 0.677 | -1 – 1.64 | 0.470 | 0.639 |
| Positive domain | 1.480 | 0.685 | 0.14 – 2.82 | 2.161 | 0.031 |
| Naltrexone | 1.353 | 0.828 | -0.26 – 2.97 | 1.634 | 0.103 |
| Curiosity rating | 0.806 | 0.116 | 0.58 – 1.03 | 6.938 | <0.0001 |
| Positive domain: Naltrexone | -1.532 | 0.852 | -3.20 – 0.13 | -1.798 | 0.073 |
| Positive domain:Curiosity | -0.292 | 0.122 | -0.53 – -0.05 | -2.390 | 0.017 |
| Naltrexone:Curiosity | -0.366 | 0.149 | -0.66 – -0.08 | -2.466 | 0.014 |
| Positive domain:Naltrexone:Curiosity | 0.393 | 0.157 | 0.09 – 0.70 | 2.507 | 0.012 |
| Random effects | | | | | |
| | Variance | sd | | | |
| Subject | 0.389 | 0.624 | | | |
| Model fit | | | | | |
| R2 | Marginal | Conditional | | | |
| | 0.36 | 0.51 | | | |
